# AltumAge: A Pan-Tissue DNA-Methylation Epigenetic Clock Based on Deep Learning

**DOI:** 10.1101/2021.06.01.446559

**Authors:** Lucas Paulo de Lima Camillo, Louis R Lapierre, Ritambhara Singh

## Abstract

Several age predictors based on DNA methylation, dubbed epigenetic clocks, have been created in recent years, with the vast majority based on regularized linear regression. This study explores the improvement in the performance and interpretation of epigenetic clocks using deep learning. First, we gathered 143 publicly available data sets from several human tissues to develop AltumAge, a neural network framework that is a highly accurate and precise age predictor. Compared to ElasticNet, AltumAge performs better for within-data set and cross-data set age prediction, being particularly more generalizable in older ages and new tissue types. We then used deep learning interpretation methods to learn which methylation sites contributed to the final model predictions. We observe that while most important CpG sites are linearly related to age, some highly-interacting CpG sites can influence the relevance of such relationships. Using chromatin annotations, we show that the CpG sites with the highest contribution to the model predictions were related to gene regulatory regions in the genome, including proximity to CTCF binding sites. We also found age-related KEGG pathways for genes containing these CpG sites. Lastly, we perform downstream analyses of AltumAge to explore its applicability and compare its age acceleration with Horvath’s model. We show that our neural network approach predicts higher age acceleration for tumors and for cells that exhibit age-related changes *in vitro*, such as replicative senescence and mitochondrial dysfunction. Altogether, our neural network approach provides significant improvement and flexibility to current epigenetic clocks for both performance and model interpretability.

---

One of the leading challenges in the field of aging research is measuring age accurately. Monitoring healthy individuals for decades to assess whether an intervention affects the aging process is prohibitive in terms of time and funding. The creation of epigenetic clocks, age predictors that use DNA methylation data, has given researchers tools to measure the aging process quantitatively. Moreover, recent works have demonstrated the effectiveness of precise epigenetic editing based on CRISPR with targeted DNA methylation or demethylation [1]. Consequently, epigenetic clocks have the potential of not only measuring aging but also guiding epigenetic interventions.

Notably, two of the most well-known predictors are the ones developed by Hannum *et al.* [2] and Horvath [3] in 2013. Hannum *et al.* developed a blood-based epigenetic clock with 71 CpG sites [2]. Then Horvath showed epigenetic clocks could also accurately predict age across tissues, developing a predictor with 353 CpG sites [3]. Horvath’s model has been widely used as it is seen as the state-of-the-art pan-tissue epigenetic clock for humans [4–7]. Both of these works used simple regularized linear regression (ElasticNet) for feature selection and prediction [8]. More recent epigenetic clocks that predict mortality also use a linear combination of features [9, 10]. ElasticNet has been widely used to develop epigenetic clocks [2, 3, 9–13]. Nevertheless, simple linear regression can display high bias and cannot capture non-linear feature-feature interactions in the data.

Interactions among variables can be taken into account by expanding the feature space with feature multiplication. However, incorporating pairwise CpG site interactions is unfeasible given the high dimensionality of the DNA methylation data. Horvath’s model [3] selected 353 CpG sites out of total 21,368 sites. If the linear regression had taken into account all pairwise interactions, the feature space would grow to over 228 million. A large number of features is especially challenging due to the relatively low number of publicly available DNA methylation samples. Given the complexity of the epigenetic regulatory network, it is likely that important interactions among CpG sites are not captured in the current epigenetic clocks developed thus far.

Recently, Galkin *et al.* [14] showed that a deep neural network model, DeepMAge, gave slightly better prediction performance than Horvath’s model in blood samples. However, the authors compared Horvath’s pan-tissue predictor to a model trained only in blood DNA methylation data. Moreover, there was no in-depth exploration of why their deep learning model outperformed the ElasticNet model. Similarly, Levy *et al.* [15] developed a deep learning framework to work with DNA methylation data that encodes the CpG sites into latent features for downstream analysis. They showed encouraging results for age prediction using a multi-layer perceptron; however, they investigated only one data set obtained from white blood cells. Therefore, currently, our understanding of the advantages of neural networks for this task in a pan-tissue setting is limited.

We introduce AltumAge, a deep neural network that uses beta values from 20,318 CpG sites common to the Illumina 27k, 450k and EPIC arrays for pan-tissue age prediction (summarized in Figure 1a). We hypothesized that a neural network using all available CpG sites would be better suited to predict pan-tissue age using DNA methylation data due to their ability to (1) capture higher-order feature interactions and (2) leverage important information contained in the thousands of CpG sites not selected by ElasticNet models. AltumAge uses multi-layer perceptron layers (similar to [14, 15]) that account for non-linear interactions by combining multiple features into each node of the network. We trained AltumAge on samples from 143 different experiments, which, to our knowledge, is the largest compilation of DNA methylation data sets for human age prediction. The publicly available data were obtained from multiple studies that used Illumina 27k and Illumina 450k arrays.

**Figure 1:**
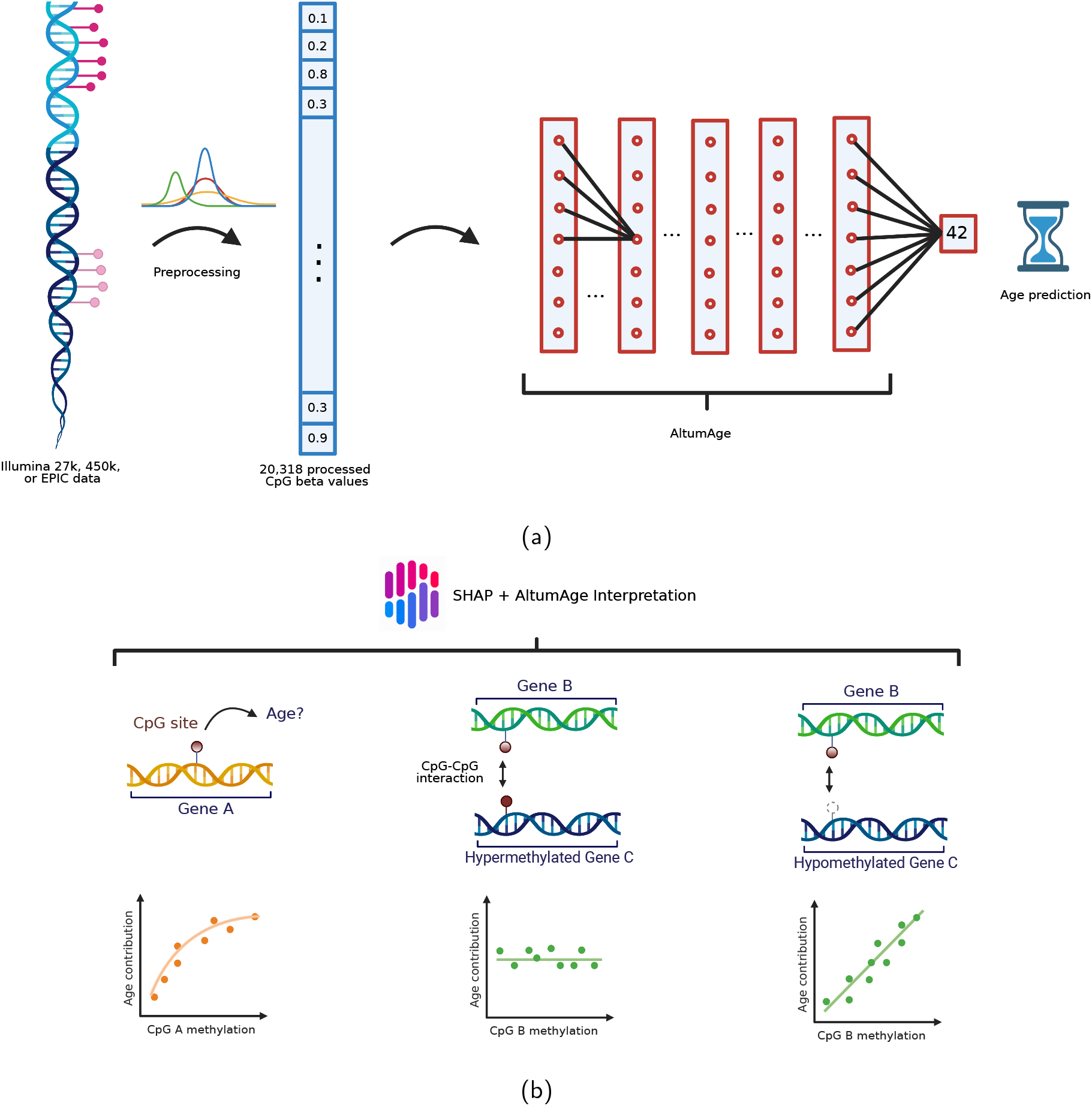
AltumAge model and interpretation. (a) DNA methylation data from Illumina 27k, 450k, or EPIC arrays are normalized with BMIQ and scaled. Then 20,318 CpG sites are selected as the input of the model. The information is processed through five hidden layers with 32 nodes each, and the values of the last hidden layer nodes are combined into a single node as the age output in years. (b) For interpretation, a Shapley-values-based method, called SHAP [16], is used to determine how the methylation status of a specific CpG site affects the age output of AltumAge. Relevant CpG sites generally present a primarily linear relationship (left) with the predicted age. However, interacting CpG sites can change such relationships. In some instances, we find that when a secondary CpG site is hypermethylated (middle), the methylation status of the first CpG is irrelevant for age prediction; when it is hypomethylated (right), then the methylation status becomes essential. Images created with Biorender.com.

We show that AltumAge has a significantly lower error for within-data set age prediction, is better able to generalize to new tissue types and older ages for cross-data set settings, and is more resistant to noise than ElasticNet. For inference, we apply the Shapley-value-based interpretation method, called SHAP [16], on AltumAge to determine the contributions of different CpG sites towards age prediction (summarized in Figure 1b). We confirm that the most important CpG sites have complex interactions resulting in non-linear relationships when predicting age. Such interactions may lead to mechanistic hypotheses on how the epigenetic network interacts to drive the aging phenotype. Additionally, we find that the most important CpG sites are proximal to CTCF binding sites. However, CpG sites in known age-related pathways (SIRT, mTOR, and AMPK) do not seem relevant for age prediction. Finally, our downstream analysis reveals that AltumAge predicts higher age for tumors, senescent cells, cells with mitochondrial dysfunction, and cells with high passage numbers than controls demonstrating its usefulness for age-related studies. Overall, we show that deep learning can improve both the performance and interpretation over the widely used ElasticNet-based models and present AltumAge as a useful tool for age prediction.

## Results

### Performance Evaluation

#### Model selection

Neural networks can capture complex variable interactions when provided with a large number of high-dimensional datasets. We hypothesized that the same would be true for age prediction with DNA methylation data. For comparison, several machine learning models were trained and validated (see Methods and Supplementary Table S1) to pick the best performing models according to median absolute error (MAE) and mean squared error (MSE). Our primary baseline method is the ElasticNet model, a linear model with implementation following Horvath [3]. We also included two traditional machine learning methods that capture non-linear relationships in the data, random forest and support vector regression; however, they performed poorly on the validation set (MAE = 6.833 and 14.229, respectively). In addition, we tried the neural network TabNet, an attentive interpretable tabular learning method [17], but the error (MAE = 4.172) was slightly worse than the baseline ElasticNet (MAE = 3.674). Lastly, we tuned the hyperparameters of the neural networks based on recent findings that highly regularized deep learning methods excel in tabular data prediction [18]. The best neural network model based on both the MAE and mean MSE was dubbed AltumAge (from the Latin *altum*, meaning “deep”). For clarity, a regularized linear regression model with Horvath’s age transformation, trained on our 143 data sets, and using the built-in hyperparameter tuning from the Python `glmnet` will be referred to as ElasticNet. On the other hand, the application of Horvath’s original 2013 epigenetic clock, originally trained on 39 data sets in that paper [3], will be referred to as Horvath’s model.

#### AltumAge outperforms linear models for within-data set age prediction

Differences in performance among epigenetic clocks can generally be explained by three factors: the DNA methylation data, the model, and the input CpG sites (or the features). For each of the 143 data sets, we split the total samples −60% for training and 40% for testing - to avoid introducing any bias in the age, gender, and tissue type distributions. The details of the data sets can be found in the Supplementary Material. We used the same training and test sets for each model to control for the data. Main results are shown in Table 1 and Supplementary Table 3.

**Table 1:** Evaluation metrics of AltumAge and different linear models in the test set. The median absolute error (MAE) and the median error are in units of year, while the mean squared error (MSE) is in units of year-squared. Also shown is the number of CpGs whose model importance is greater than zero.

| Model | CpGs | MAE | MSE | R | Median Error |
| --- | --- | --- | --- | --- | --- |
| AltumAge | 20318 | **2.179** | **31.324** | **0.979** | -0.153 |
| AltumAge with ElasticNet CpGs | 1218 | 2.328 | 32.005 | 0.978 | 0.093 |
| AltumAge with Horvath's CpGs | 353 | 2.398 | 32.559 | 0.978 | **0.015** |
| ElasticNet | 1218 | 2.773 | 40.340 | 0.973 | -0.041 |
| Linear Regression with Horvath's CpGs | 353 | 3.247 | 51.628 | 0.965 | -0.025 |
| Horvath's model | 353 | 3.659 | 74.161 | 0.949 | 0.186 |

We hypothesized that our large and diverse DNA methylation data might improve performance compared to other epigenetic clocks irrespective of model type, adding a confounding variable to any performance improvement seen with AltumAge. To understand the magnitude of such effect, we compared a replication of Horvath’s model as seen in [3] with a linear regression trained on our 143 data sets using the same set of 353 CpG sites. Indeed, the regression trained with our data has a lower error (MAE = 3.247 vs. 3.659; MSE = 51.628 vs. 74.161). ElasticNet, with its selected 1218 CpG sites trained with our data, further improves the performance (MAE = 2.773, MSE = 40.340). This result shows that a larger training data set helps the age prediction performance.

Next, we aimed to determine whether the model type, i.e., a linear regression vs. a neural network, would significantly impact the performance. We, therefore, compared the aforementioned linear models with the neural network AltumAge using the same set of features. AltumAge outperformed the respective linear model with Horvath’s 353 CpG sites (MAE = 2.398 vs. 3.247, MSE = 32.559 vs. 51.628) and ElasticNet-selected 1218 CpG sites (MAE = 2.328 vs. 2.773, MSE = 32.005 vs. 40.340). This result shows that AltumAge outperforms linear models given the same training data and set of features.

Lastly, to compare the effect of the different sets of CpG sites, we trained AltumAge with all 20,318 CpG sites available and compared the results from the smaller sets of CpG sites obtained above. There is a gradual improvement in performance for AltumAge by expanding the feature set from Horvath’s 353 sites (MAE = 2.398, MSE = 32.559) to 1218 ElasticNet-selected CpG sites (MAE = 2.328, MSE = 32.005) to all 20,318 CpG sites (MAE = 2.179, MSE = 31.324). This result suggests that the expanded feature set helps improve the performance, likely because relevant information in the epigenome is not entirely captured by the CpG sites selected by an ElasticNet model.

Overall, these results indicate that even though more data samples lower the prediction error, AltumAge’s performance improvement is greater than the increased data effect.

A direct comparison of AltumAge and Horvath’s model reveals that AltumAge has fewer tissue types with a high MAE. In his 2013 paper, Horvath noticed poor calibration of his model in breast, uterine endometrium, dermal fibroblasts, skeletal muscle, and heart [3]. In our test data, a similarly poor predictive power was found for these tissue types for Horvath’s model (breast MAE = 9.462; uterus MAE = 5.798; fibroblast MAE = 10.863; muscle MAE = 9.525; heart not included). AltumAge, on the other hand, had much lower errors for them (MAE = 3.872, 3.049, 5.757, 2.512 respectively). Furthermore, Horvath’s model had an MAE > 10 years in 22 tissue types in the test data. AltumAge, on the other hand, had an MAE > 10 in only three tissue types.

Supplementary Figure S1 in particular shows how AltumAge, in contrast to Horvath’s model, does not underestimate older ages ( > 60 years) to such an extent (median error = −2.542 vs. −4.651). Better performance in older age is fundamental in defining biomarkers of age-related diseases of which age is the biggest risk factor. Horvath’s model tends to underestimate such population partly due to CpG saturation (beta value approaching 0 or 1 in certain genomic loci) [19]. Another reason might be the assumption that age-related CpG changes are linearly correlated with age after 20 years of age. AltumAge resolves these two problems by incorporating an expanded feature set and not using any pre-defined age transformation function that might inject bias in the data processing.

Of note, we were unable to compare AltumAge with DeepMAge [14], another deep learning framework. Unfortunately, neither the code for DeepMAge nor a complete description of its architecture is available.

#### AltumAge is more generalizable than ElasticNet in older ages and in non-blood tissue types

Leave-one-data-set-out cross-validation (LOOCV) provides a way to understand the generalization potential of a model to new unseen data sets. We performed this LOOCV analysis by leaving out the training samples of each data set (out of the 143) during model fitting. Therefore, the model training was performed using the training set of 142 data sets. Next, we evaluated the performance of this model on the test set of the left-out data set. Consequently, we trained 143 different models in total to evaluate the LOOCV performance for all 143 data sets (Figure 2, Table 2).

**Figure 2:**
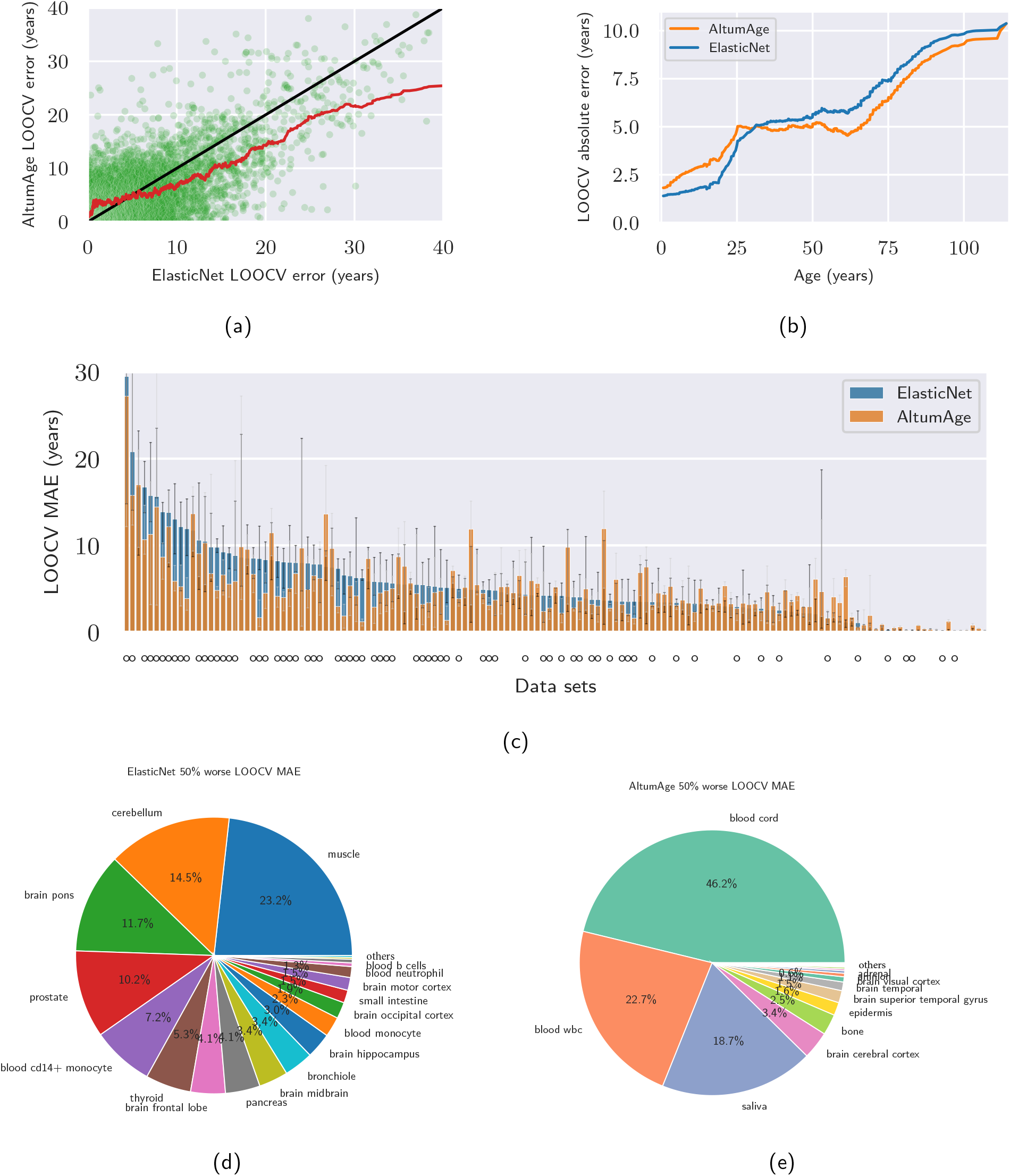
Comparison of leave-one-data-set-out cross-validation (LOOCV) performance between AltumAge and ElasticNet. (a) Scatter plot contrasting the LOOCV absolute error of each model by sample. The black line separates the region in the graph in which AltumAge performs better (bottom right) versus where ElasticNet is superior (top left), and the red line is a 100-sample rolling mean. AltumAge outperforms ElasticNet, particularly in difficult-to-predict tissue types. (b) The 1000-sample rolling mean of the LOOCV absolute error of each model by age. AltumAge has a lower absolute error for age >30 years on average. (c) Bar plot showing the LOOCV median absolute error (MAE) by data set for each model, with 95% confidence interval error bars calculated from 1000 bootstrap iterations. A circle below a bar represents data sets in which AltumAge had a lower LOOCV MAE than ElasticNet. (d) Pie plot showing which tissue types ElasticNet had at least a 50% worse MAE than AltumAge. (e) Pie plot with the converse. Overall, AltumAge can better generalize to more tissue types, whereas most of the improved ElasticNet performance comes from blood-based tissues.

**Table 2:** Leave-one-data-set-out cross validation evaluation metrics for AltumAge and ElasticNet. The median absolute error (MAE) and the median error are in units of year, while the mean squared error (MSE) is in units of year-squared.

| Model | MAE | MSE | R | Median Error |
| --- | --- | --- | --- | --- |
| AltumAge | **3.578** | **67.329** | **0.955** | **-0.130** |
| ElasticNet | 3.641 | 72.317 | 0.951 | -0.187 |

Since AltumAge uses 20,318 CpG sites, we expected it to be more prone to noise and overfitting than a model with low variance such as ElasticNet, which effectively uses only a subset of CpG sites. Nevertheless, we see that AltumAge performs better than ElasticNet in both MSE (67.329 vs. 72.317) and MAE (MAE = 3.578 vs. 3.641, Wilcoxon signed-rank test p = 0.0036).

Next, we analyzed the results stratified by absolute error, age, data set, and tissue type. Figure 2a shows a scatter plot of the LOOCV absolute error for each sample according to AltumAge and ElasticNet. Points above the black line favor ElasticNet while the opposite favors AltumAge. As shown by the 100-window rolling mean line, for samples with an absolute prediction error > 3.341 years, on average, AltumAge performs better. This observation is particularly apparent for large deviations. This result indicates, alongside the lower MSE, that AltumAge is more resistant to outliers than ElasticNet when generalizing to new samples.

Stratifying the results by age can give insights into particular strengths and weaknesses of each model. For example, while both models capture the age-related epigenetic drift given the correlation between absolute error and age (AltumAge Pearson’s R = 0.376; ElasticNet Pearson’s R = 0.423), AltumAge performs better on average for samples with age >30 years (Figure 2b). This result suggests that AltumAge better captures epigenetic changes during aging while ElasticNet better understands the developmental epigenome, since epigenetic changes during childhood and puberty are related to development but after they are mostly due to aging [20].

Finally, we analyzed the performance of each model by data set and tissue type. As shown in Figure 2c, AltumAge performs better than ElasticNet in the LOOCV MAE for data sets with a higher error in performance, i.e., that are more challenging to predict. In comparison, ElasticNet is superior for the ones with a lower error. We observe that most data sets with a low MAE are from newborns or blood samples, and the training set is skewed towards blood-based samples (see Supplementary Figure S2). Therefore, we hypothesized that ElasticNet may be simply performing better for overrepresented tissue types in the training set. To check this, we looked at tissue types for which ElasticNet had a 50% worse LOOCV MAE than AltumAge (capturing a large deviation) and vice versa. As expected, ElasticNet does not generalize as well to a large variety of tissue types (Figure 2d). At the same time, it performs better in blood-based samples (Figure 2e). These observations imply that AltumAge can better generalize to more tissue types, likely capturing global age-related epigenetic patterns, while ElasticNet could be focusing primarily on blood changes.

As each model has its benefits and drawbacks, we checked the performance of an ensemble of both methods. Interestingly, we observe a substantial decrease in both MAE (3.269) and MSE (61.962) by averaging the predictions of both models. These results indicate that combining deep learning and linear model predictions may further improve the age prediction performance.

#### AltumAge is more robust to noise than ElasticNet

Another desired property of epigenetic clocks is reliability. Noise derived from the experimental procedure, biological or technical replicates may negatively influence the model’s reliability. AltumAge was trained with Gaussian noise and adversarial regularization to be more robust against random variation [21]. Gaussian noise introduces normally distributed fluctuations in between hidden layers. Adversarial regularization includes artificial observations with subtle modifications in the loss function that attempt to fool the model into increasing the error. To assess the robustness of AltumAge and ElasticNet to noise, we gradually added artificial Gaussian noise in the beta value of each CpG site up to one standard deviation in the within-data test set and tracked MAE (Figure 3a) and MSE (Figures 3b). As expected, the error grows much faster in the ElasticNet model, particularly with the MSE, which is more swayed by outliers.

**Figure 3:**
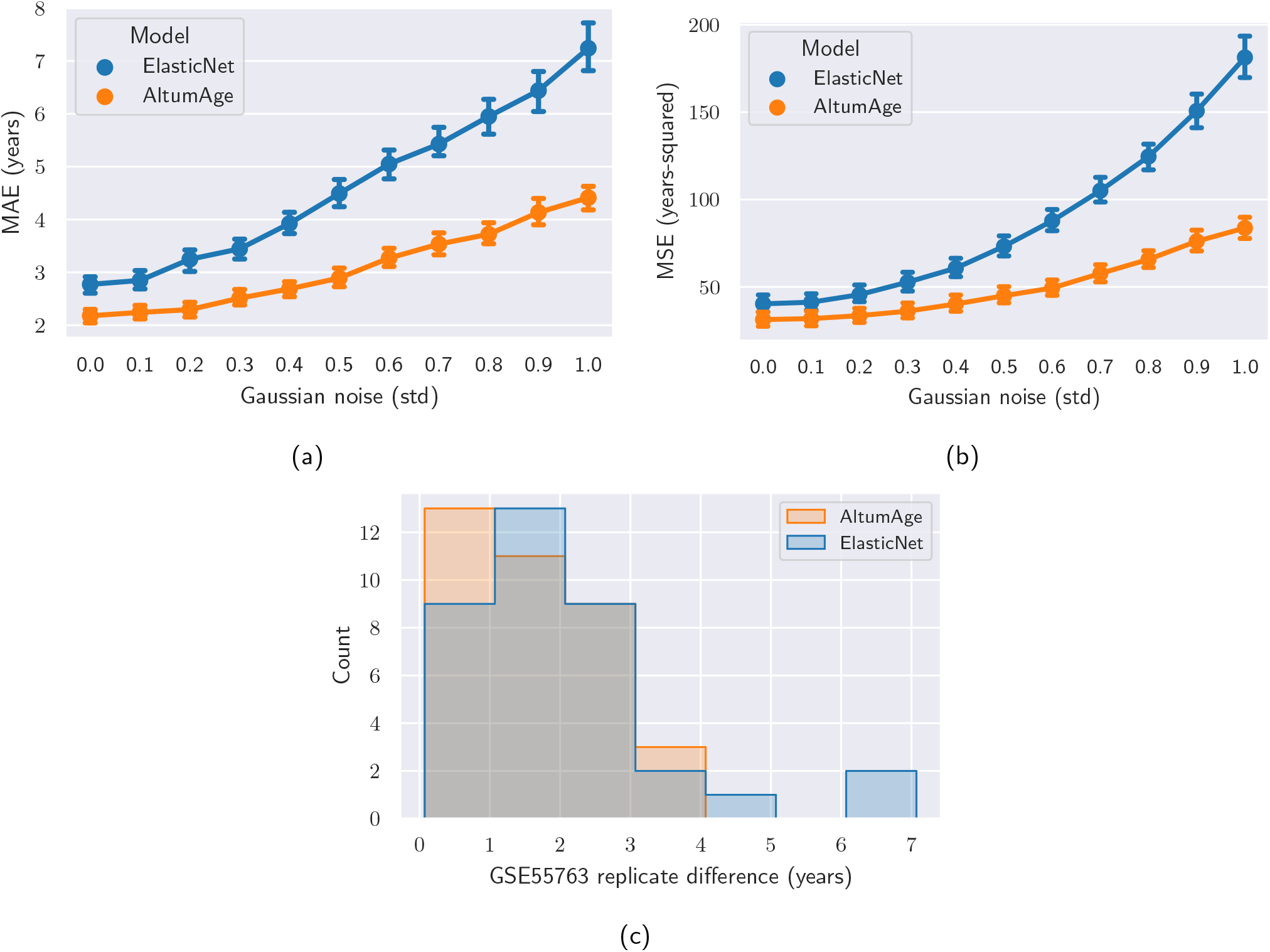
Comparison of resistance to noise between AltumAge and ElasticNet. (a, b) Point plots show the increase in median absolute error (MAE) and mean squared error (MSE) per model when adding artificial Gaussian noise of up to one standard deviation for each feature. AltumAge is more resistant to noise in both metrics. Shown are the 99% confidence interval error bars calculated from 1000 bootstrap iterations. (c) Histogram of the difference in predicted age between two technical replicates in an independent whole blood data set (GSE55763). AltumAge has a lower median and maximum deviations than ElasticNet.

Furthermore, we examined a new independent whole blood data set GSE55763 (not used in training or testing), which contains 2 technical replicates for each of its 36 samples. Ideally, the difference in prediction between the replicates would be zero. As shown in the histogram in Figure 3c, the median absolute difference for AltumAge is 1.516 years, whereas for ElasticNet, 1.845 years, while the maximum absolute difference is 3.597 and 6.654 years, respectively. Despite no significant difference in distributions — likely due to the small sample size — the models differ in whether they capture an artifact effect between replicates. As anticipated, we do not observe a statistically significant effect from replicate one to two for AltumAge (linear mixed-effects p = 0.441). However, we see that ElasticNet predicted a slightly higher age of 0.848 years for replicate two (linear 5 mixed-effects p = 0.031). Overall, the results highlight that resistance to random noise may help in real-world 6 scenarios, increasing model robustness and reliability.

### Inference

Neural networks, particularly in the context of deep learning, used to be seen as “black-box” methods, as their interpretability was difficult. On the other hand, regardless of the predictive power of ElasticNet models, they are easily understandable. Recently, various methods have been proposed to extract the contribution of features towards prediction in neural networks. They include interpretation based on model gradients [22–24], attention [25], among others. One such inference method is SHAP [16], which uses a game-theoretic approach to aid in the explanation of machine learning methods. It can measure how one feature contributes to the output of deep neural networks. For our case, the SHAP value can be conceived as how much the value of one CpG site affects the age output of the model in years. Through the architecture of neural networks, it can also determine which CpG sites most highly interact with each other.

We present results for model inference using SHAP that assist with understanding AltumAge. To support the results obtained by SHAP, we also applied another method of determining feature importance called DeepPINK [26] (see Supplementary Information).

#### AltumAge captures relevant age-related CpG-CpG interactions

Epigenetic modifications can significantly influence gene expression. They can also impact genes that affect other epigenetic changes. Therefore, some CpG sites interact with others through the gene expression network and can work in tandem. Through SHAP, we show that AltumAge can measure how hyper- or hypomethylation of secondary CpG sites affects the relationship of a CpG of interest and age. Supplementary Figure S7 shows scatter plots of the nine most important CpG sites based on SHAP-based importance values assigned to the CpG sites of the samples in the test set. These nine CpG sites account for 1.99% of the total model importance (Supplementary Figure S3). The dependence plots show both the relationship of a CpG site with the predicted age and how that relationship can be affected by the value of a secondary CpG site for a DNA methylation sample. This secondary CpG site has the highest interaction with the CpG of interest, as determined by SHAP values. One way to understand the effect of the secondary CpG site is to focus on the samples in the top and bottom deciles of its methylation value, looking for any differences that may arise due to hyper- or hypomethylation respectively. We uncovered three different types of relationships between CpG site methylation value and age: (1) completely linear, which are independent of CpG-CpG interactions; (2) bivalently linear, whose slope is dependent on a secondary CpG site; and (3) non-linear, affected by a secondary CpG site.

Out of the top nine CpG sites, only cg04084157 (Figure 4a), the fifth most important, shows a completely linear relationship. We subdivided the samples in the test set into the top and bottom deciles for cg07388493, the most highly interacting CpG site. Both subsets display linear relationships (Figure 4b) with indistinguishable regression coefficients (Z-test^1^ p = 0.346). Consequently, we observe that the effect of cg04084157 on the age output is independent of the value of the most highly interacting CpG site, cg07388493.

**Figure 4:**
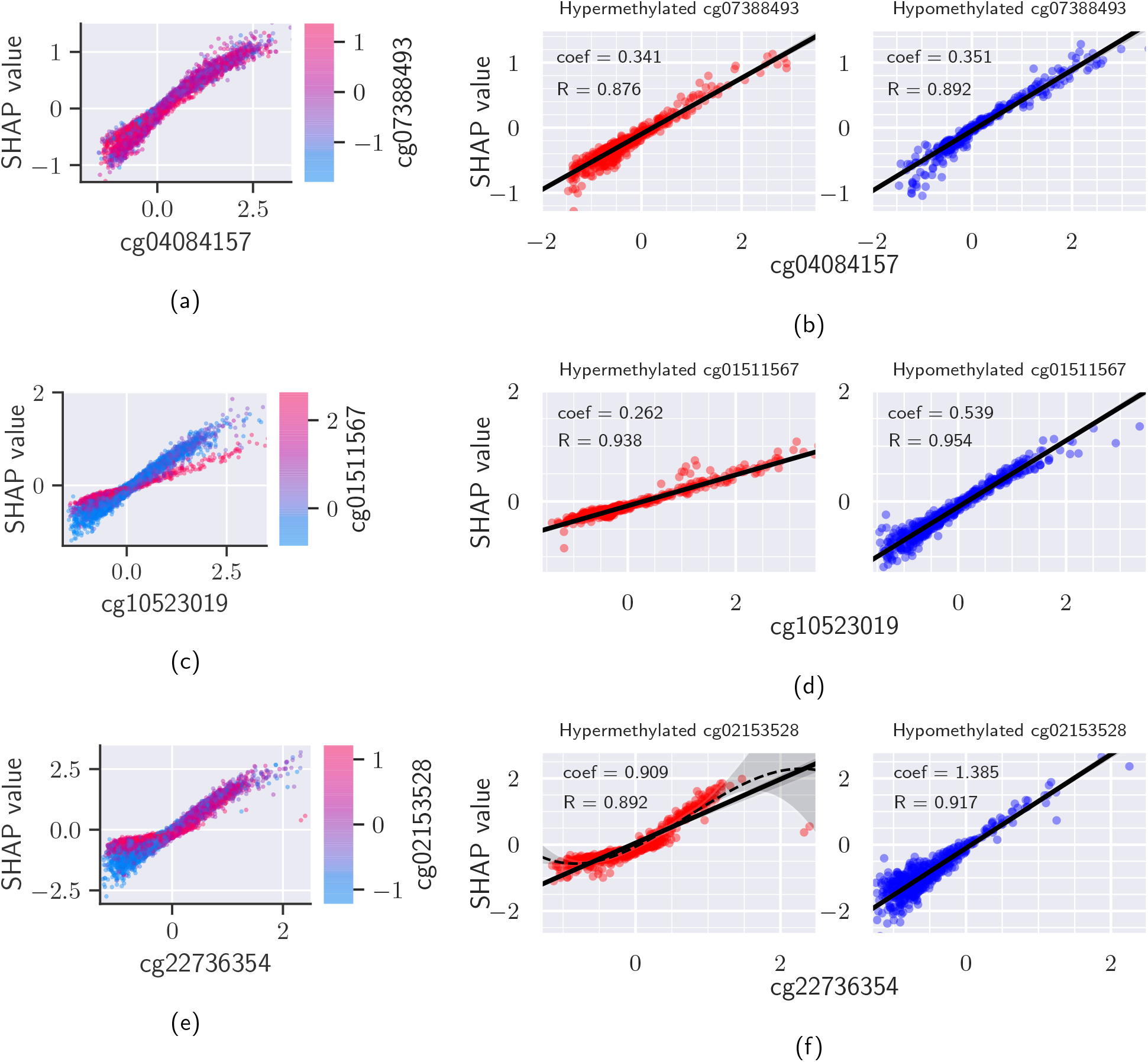
Three main types of relationship between the scaled beta value of a CpG site and age according to SHAP. Dependence plots of the fifth (a), seventh (c), and first (e) most important CpG sites exemplify the three types of relationship. Samples into the top (red) and bottom (blue) deciles of the most highly interacting CpG site were divided, representing hyper- and hypomethylation respectively. The relationshps are completely linear (b), bivalently linear (d), and non-linear (f). Regression lines are shown in (b), (d), and (f) with a 95% confidence interval calculated from 1000 bootstrap iterations. A cubic regression is also shown in (f) to demonstrate the better fit of the non-linear model.

The effect of cg10523019 (Figure 4c), the seventh most important CpG site, displays a bivalently linear relationship. The regression coefficient when cg01511567, the most highly interacting CpG site, is hypomethylated (coef = 0.539) is more than twice when it is hypermethylated (coef = 0.262, Z-test p = 6.3e-238, Figure 4d). This dual response may also shine a light on relevant age-related biological processes. cg01511567 is located in the gene SSRP1, a subunit of the chromatin transcriptional elongation factor FACT, and cg10523019 lies in RHBDD1, a gene involved in proteolysis and apoptosis. Since RHBDD1 has a long transcript of 163,280 base pairs (about seven times the median gene length [27]), FACT likely aids RHBDD1’s transcription. When SSRP1 is expressed (with cg01511567 hypomethylated), RHBDD1 has more influence on age prediction. However, if SSRP1 is repressed (with cg01511567 hypermethylated), the methylation status of cg10523019 becomes less relevant as transcription of RHBDD1 may become deficient due to lack of SSRP1. Laboratory experiments would have to be performed to more thoroughly characterize these relationships; however, it is possible to obtain data-driven hypotheses from these dependence plots.

An example of a non-linear CpG-age relationship comes from the top most important CpG cg22736354 (Figure 4e). When cg02153528, the most highly interacting CpG site is hypomethylated, the relationship between cg22736354 and age output is still linear, despite its heteroscedasticity (or unequal scatter). However, when it is hypermethylated, it becomes non-linear (Figure 4f). While Pearson’s correlation coefficient for a straight line is high (0.892), the residual plot (Supplementary Figure S8) shows underestimation at the boundaries with overestimation in the center. This pattern demonstrates non-linearity; a cubic regression corrects the bias of under and overestimation and increases the correlation coefficient to 0.947. Overall, our results using SHAP values demonstrate that AltumAge captures the non-linear interaction between CpG sites.

Note that despite their important effects on the predicted age, some of the CpG sites that interact with the most important CpG sites may themselves not be particularly relevant for the output. For example, the aforementioned cg02153528, the CpG with the highest interaction with the most important CpG site, ranks 11052 and 14928 according to SHAP and DeepPINK, respectively, out of 20,318. These results suggest that ElasticNet may miss DNA loci that regulate other loci in aging, and this may partly explain AltumAge’s performance improvement compared to ElasticNet.

#### Characterization of CpG sites by model interpretation

CCCTC-Binding factor (CTCF) is a transcription factor involved in the negative regulation of several cellular processes. It also contributes to long-range DNA interactions by affecting chromatin architecture. We examined whether CpG sites with a higher SHAP importance were closer to CTCF binding sites. The 353 important CpG sites selected by Horvath’s model were not closer to the CTCF binding sites when compared to the 21,368 control CpG sites from which the paper’s ElasticNet model was trained (Mann-Whitney U-test p-value = 0.991). As for AltumAge, since it uses all of the 20,318 CpG sites as features, we compared the top 1000 CpG sites to the control, as the ElasticNet model applied on the full data set selects 1218 sites as important. These sites comprise 50.8% of SHAP importance. In line with previous studies [28–30], we find that the selected important CpG sites are overwhelmingly closer to CTCF binding sites (Mann-Whitney U-test p-value = 0.00636, Supplementary Figure S4). This observation suggests that epigenetic alterations proximal to such loci that are involved in chromatin packing by affecting CTCF binding may be captured by AltumAge. This result is relevant because chromatin structure modifications have been associated with aging (see review [31]).

Due to the close relationship between chromatin and aging, we hypothesized that different chromatin states would influence the importance of each CpG site. ChromHMM is a Hidden Markov Model used for the characterization of chromatin states [32]. Annotations for several cell lines and tissue types are widely available online. Since AltumAge is a pan-tissue epigenetic clock, we used the mode of the 18-state annotation from 41 different tissues obtained from ENCODE for each CpG location [33] (Supplementary Figure S5, Supplementary Table S2). CpG SHAP importance values are indeed impacted by ChromHMM state (Kruskal-Wallis H-test p-value = 5.756e-22). The chromatin state with the highest SHAP normalized median importance was a genic enhancer (SHAP importance = 9.45e-10%, top 69th percentile of all CpG sites). This result supports the idea that some enhancers might be highly relevant to aging [34]. Of note, the chromatin state with the highest DeepPINK normalized median importance was heterochromatin (DeepPINK importance = 2.47e-14%, top 66th percentile of all CpG sites).

#### Aging-related pathways

One of the main interpretation advantages of AltumAge compared to ElasticNet is that the former effectively uses a much larger feature space. CpG sites in aging-related genes are often not selected within the couple hundred features of an ElasticNet model, thus making analyses of these CpG sites of interest impossible. AltumAge allows a closer look at the relationship of CpG sites in aging-related pathways even when these CpG sites are not particularly important for the final age prediction. It is worth analyzing the relative importance of CpG sites in well-known age-related pathways such as SIRT, mTOR, and AMPK [35–37].

Unexpectedly, most of the CpG sites in SIRT genes do not appear relevant, at least directly, for age prediction using AltumAge. Located in SIRT2, cg27442349, accounting for 0.0389% of the total SHAP importance and ranked 574, has the highest SIRT SHAP importance value (Supplementary Figure S6). Other SIRT CpG sites were much below rank 1000.

Out of the 67 proteins participating in the mTOR signaling pathway according to the PID Pathways data set [38], cg11299964, located in MAPKAP1, has the highest SHAP importance of 0.075%, ranking 119. Surprisingly, mTOR was not particularly relevant, with its most important CpG site being cg07029998 (SHAP importance = 0.022%, rank 1544) (Supplementary Figure S6).

In terms of the AMPK pathway, out of CpG sites in genes for the proteins that directly activate or inhibit AMPK from the KEGG database [39], cg22461835, located in ADRA1A, has the highest SHAP importance of 0.093%, ranking 69 (Supplementary Figure S6). Most, however, ranked below 1000.

We also performed KEGG pathway analysis on the genes related to the top-ranking nine CpG sites using KEGGMapper [40]. We found the following genes associated with four of them - NHLRC1, involved in proteolysis; EDARADD, involved in NF-kappa B signaling pathway; NDUFS5, involved in metabolic pathways, including oxidative phosphorylation and thermogenesis; and FZD9, involved in a range of age-related diseases, including cancer and neurodegeneration. Note that DNA methylation affects gene expression depending on its position. A methylated CpG site in an enhancer, promoter, or gene body may impact gene regulation differently. These findings show how methylation in specific loci in aging-related pathways can contribute to age prediction. This insight may not be possible to obtain using ElasticNet due to its focus on selecting only the most important CpG sites related to aging. For example, only cg11299964 (from MAPKAP1 mentioned above) was present among the 353 sites selected by Horvath’s model.

### Potential biological applications

The age acceleration, defined as the predicted age minus the real age, of epigenetic clocks have been shown to be related to several biologically relevant events and characteristics, such as obesity [41], menopause [42], diet [43], heart disease [44], anxiety [45], and even socioeconomic status [46], among others. Given the observed performance of AltumAge, we explore its applicability to such studies, for which Horvath’s model has been a popular choice. Therefore, we assess AltumAge’s age acceleration performance with respect to the Horvath’s model in the following sections to understand the usefulness of its age prediction for downstream analyses.

#### AltumAge differentiates cells with age-related hallmarks

To assess whether AltumAge can capture biologically relevant changes, we examined independent data sets (not used for training or testing) to understand the effect of cellular senescence, cell passage number, mitochondrial depletion, NLRP7 knockdown, and transient cellular reprogramming. We compared AltumAge with Horvath’s model as a benchmark.

Cellular senescence is a well-known hallmark of aging. A study (with accession ID GSE91069 [47]) provides methylation data from cultured fibroblasts in early passage (EP), near senescence (NS), oncogene-induced senescence (OIS), and replicative senescence (RS). Despite the small sample size (n = 15), AltumAge displays a sequential increase in median predicted age going from EP to NS to OIS to RS (Figure 5a). We observe an increase between EP and RS (one-sided Mann-Whitney U-test p-value = 0.047). Horvath’s model also captures a sequential increase from EP to NS to RS, although with lower confidence in differentiating EP from RS (one-sided Mann-Whitney U-test p-value = 0.131).

**Figure 5:**
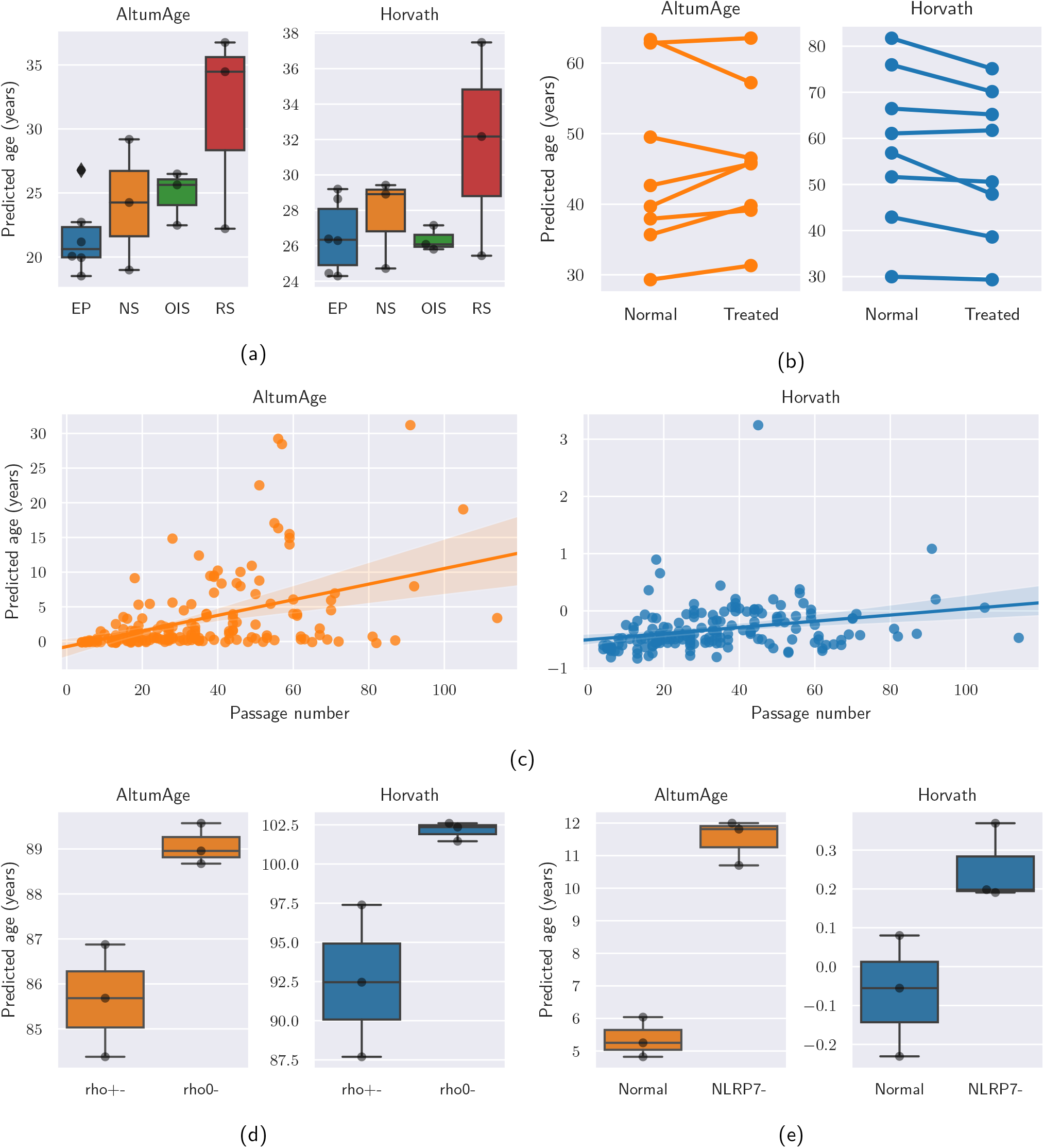
Analysis of the effect of cellular senescence, cell passage number, mitochondrial depletion, NLRP7 knockdown, and transient cellular reprogramming on predicted age comparing AltumAge and Horvath’s model. (a) Box plots showing predicted age for cultured fibroblasts in early passage (EP), near senescence (NS), oncogene-induced senescence (OIS), and replicative senescence (RS). (b) Point plot showing age prediction in human fibroblasts and endothelial cells before and after transient reprogramming. (c) Scatter plot of predicted age of iPSCs and ESCs by passage number with best fit line with 95% confidence interval calculated from 1000 bootstrap iterations. (d) Box plots showing predicted age in 143B cells with mitochondrial depletion (rho0-) or control (rho+-). (e) Box plots showing predicted age of H9 ESCs with NLRP7 knockdown (NLRP7-) or control (Normal).

Given that AltumAge detects a tentative link between replicative senescence and higher age, we looked into a study (with accession ID GSE30653 [48]) which contains information of induced pluripotent stem cells (iPSCs) and embryonic stem cells (ESCs) by passage number. We observe that AltumAge detects a strong correlation (Pearson’s R = 0.436, p-value = 3.187e-09) between the predicted age and the passage number. As shown by Figure 5c, cells begin with a slightly negative age that increases markedly as they are passaged. Horvath’s model also detects a significant correlation (Pearson’s R = 0.273, p-value = 3.245e-04), albeit it is weaker when compared to AltumAge. The increase in age with passage number is also much more subtle. The suggestive difference in cultured fibroblasts from EP to RS in addition to the relatively large effect of passage number on predicted age indicates that AltumAge is sensitive to replicative senescence.

Mitochondrial dysfunction is another important hallmark of aging. A study (with accession ID GSE100249 [49]) contains data on 143B cells chronically depleted of mitochondrial DNA (rho0-). Both AltumAge and Horvath’s model predict a higher age for cells with mitochondrial dysfunction (one-sided Mann-Whitney U-test p-value = 0.050, Figure 5d).

The NLRP gene family of receptors, primarily expressed in immune cells, is involved in the normal response to inflammation. Mutations in some of these genes are involved in immune system malfunction, excessive inflammation, and disease [50–52], suggesting it may also have ramifications in aging. Moreover, cells with NLRP7 knockdown display aberrant CpG methylation patterns [53]. We, therefore, analyzed DNA methylation data from H9 ESCs (accession ID GSE45727 [53]) with or without NLRP7 knockdown. AltumAge and Horvath’s model predict a higher age for knockdown cells (one-sided Mann-Whitney U-test p-value = 0.050, Figure 5e). When H9 cells are exposed to BMP4 differentiating medium, AltumAge is still able to capture the increase in age (one-sided Mann-Whitney U-test p-value = 0.050), while Horvath’s model predictions are not as confident (one-sided Mann-Whitney U-test p-value = 0.100, Supplementary Figure S9).

Lastly, we investigated whether AltumAge could capture a rejuvenation event caused by transient expression of several reprogramming factors in aged human fibroblasts and endothelial cells (GSE142439) [54]. Sarkar et al. [54] demonstrated that several biomarkers such as H3K9me3, SIRT1, HP1γ, and β-galactosidase were restored to a youthful state after treatment. Horvath’s model was able to capture a decrease in epigenetic age (linear mixed-effects model p-value = 0.004, Figure 5b). Interestingly, there was no difference before and after the intervention according to AltumAge (linear mixed-effects model p-value = 0.845, Figure 5b). While the researchers tracked the cells until six days after treatment, it is possible that the apparent restoration of youthful biomarkers would not endure. Indeed, studies have shown that transient reprogramming causes only temporary rejuvenation [55–57]. Altogether, AltumAge performs as well as Horvath’s model for most cases and, more importantly, it captures an expanded global methylation landscape. It can also robustly recognize age-related epigenetic patterns while potentially avoiding overestimating the impact of temporary interventions.

#### AltumAge predicts higher age acceleration for cancer

Cancer cells display several genetic and epigenetic aberrations which have been related to aging and mortality by epigenetic clocks [58–60]. Liu et al. [61] have reported that some age predictors consistently estimate higher age acceleration for tumors, whereas others show tissue-specific behavior. Therefore, we examined the age acceleration of cancer samples from 14 data sets comprising 10 tissue types in total for AltumAge, using Horvath’s model as a benchmark (Figure 6). Overall, Horvath’s model was not able to differentiate between normal and tumor samples (Mann-Whitney U-test p-value = 0.135, Figure 6a). Its median age acceleration for cancer was higher in five tissue types and lower in another five. AltumAge, in contrast, predicts overall higher age acceleration for cancer when compared to normal tissue by 4.832 years (Mann-Whitney U-test p-value = 1.55e-17, Figure 6b). The only two tissue types in which AltumAge estimates a slightly lower age acceleration for cancer are colon and nasopharyngeal tumors, possibly due to their smaller samples size. These results indicate that AltumAge can generally differentiate between normal and cancerous tissue by predicting a higher age acceleration and could be useful to studies focusing on relationship between cancer and aging.

**Figure 6:**
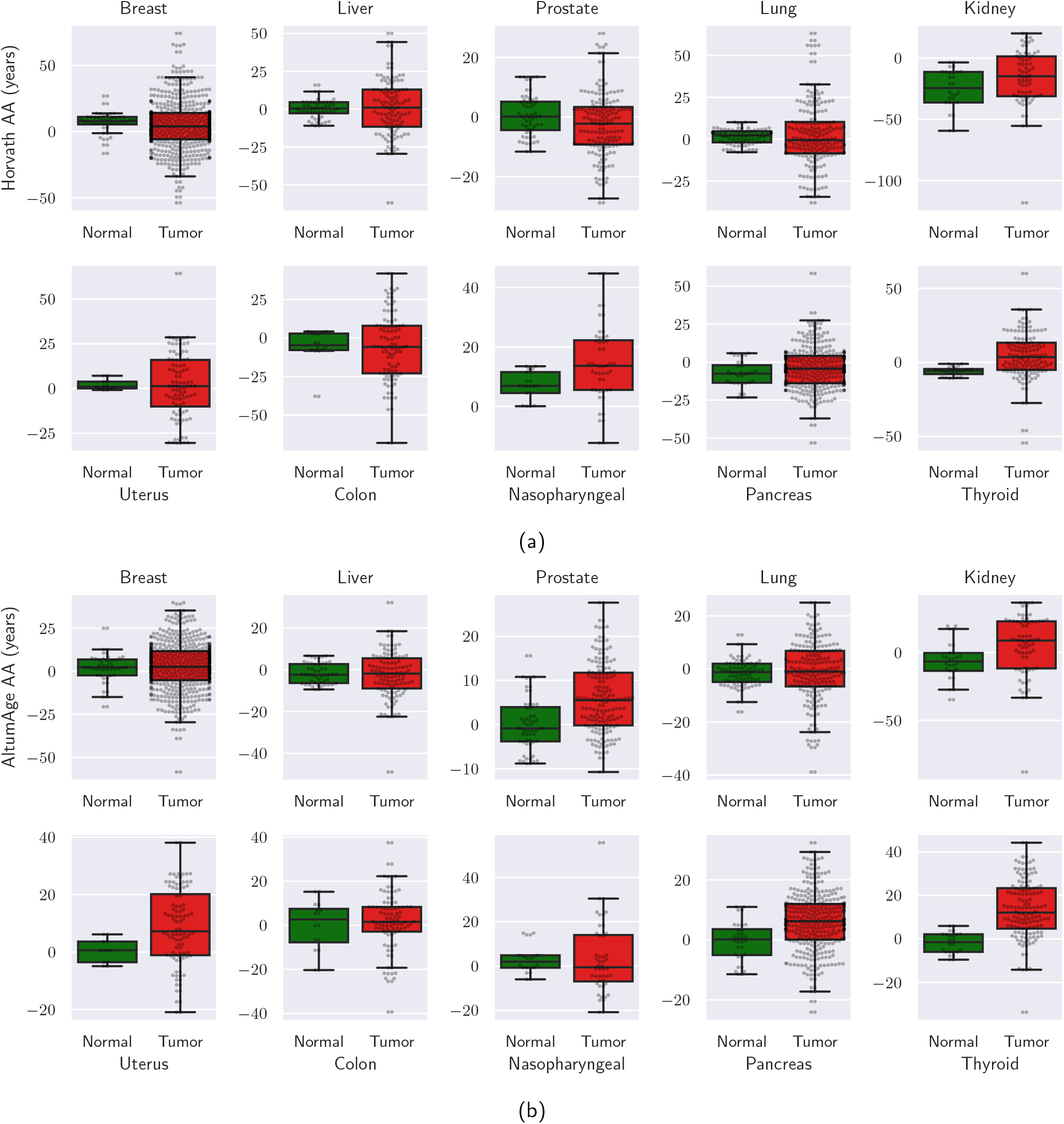
Box plots showing age acceleration (AA) in normal and cancerous samples by tissue type according to Horvath’s model (a) and AltumAge (b). Horvath’s model does not consistently predict higher AA for cancer, with tissue-specific behavior. On the other hand, AltumAge generally predicts higher AA for tumors in all but two cancer tissue types with the two lowest sample numbers (colon and nasopharyngeal).

## Discussion

The creation of new quantitative aging measurements has been rapidly expanding with the burgeoning field of the biology of aging. Epigenetic clocks are a tool that can aid researchers to understand better and to measure the aging process. In 2013, Horvath showed it was possible to use just a couple of CpG sites to predict a person’s age based on DNA methylation accurately. It was a giant leap in the field. However, his 2013 ElasticNet model or other versions of linear models are still widespread despite recent advances in machine learning. The accuracy of such linear models was so good that it was difficult to imagine a model significantly outperforming it [62]. Other deep learning methods, which slightly outperform ElasticNet, have focused thus far only in a single tissue type [14] [15].

We show that our neural network-based model, AltumAge, is an overall better age predictor than ElasticNet. While our more comprehensive and larger data does improve the performance, the capability of neural networks to detect complex CpG-CpG interactions and the expanded feature set with 20,318 CpG sites also contribute to its lower error. For within-data set prediction — which is the case for several studies which create a new epigenetic clock — AltumAge performs drastically better than state-of-the-art methods. Even for LOOCV analysis, while the improved performance of AltumAge over ElasticNet was not as substantial, it performed better in older ages and new tissue types. Arguably, a more generalizable model like AltumAge can better capture pan-tissue age-related changes.

Deep learning models have shown promise in several biological tasks, given their good performance on unstructured data. They have been for many years seen as “black-box” models, but new interpretation methods have made it possible to get interesting insights for them. Our interpretation of AltumAge provides a detailed relationship between each of the 20,318 CpG sites and age, showing that while most CpG sites are mostly linearly related with age, some important ones are not. Given recent advances in epigenetic editing [1], finding these DNA methylation sites to delay or reverse aging may be necessary for future interventions to tackle the disease. AltumAge allied with other deep learning inference methods can provide information on highly interacting CpG sites. Sometimes the primary locus of an epigenetic editing intervention, given its place in the genome, may be difficult to target because of the chromatin structure. Consequently, knowing secondary CpG sites that affect how the CpG of interest interacts with age could guide such interventions. We show that one can obtain biological hypotheses for the same from the data using AltumAge. For example, we observe that cg01511567 located inside the gene SSRP1 could regulate cg10523019, which lies in RHBDD1. Analysis of ChromHMM annotations shows that the top-ranking CpG sites are associated with gene regulatory regions and CTCF binding sites. Finally, we highlight the age-related KEGG pathways obtained for genes with these CpG sites, indicating that the model is learning valuable biological information from the data.

We also explore how age acceleration as determined by AltumAge has potentially meaningful biological applications. AltumAge predicts higher age acceleration for cells with mitochondrial dysfunction and replicative senescence, similarly to Horvath’s model. More importantly, AltumAge displays a much higher correlation between cell passage number and predicted age. Moreover, AltumAge, in contrast to Horvath’s model, predicts higher age acceleration for cancer. It seems that epigenetic clocks highly predictive of mortality show this behavior consistently [61]. Age acceleration in tumors can be thought of as a further deviation from the original state in Waddington’s landscape. Intriguingly as well, AltumAge does not uncover a rejuvenation event from transient reprogramming [54]. This observation is possibly due to the temporary nature of the rejuvenation event, which may not globally change age-related DNA methylation patterns. Since AltumAge considers a much larger portion of the epigenome, it may be more resistant to detecting momentary rejuvenation.

In future work, it would be interesting to create a deep learning model with Illumina’s EPIC array with the roughly 850 thousand CpG sites to understand more deeply how genomic location can affect influence in aging. In addition, by having several CpG sites in a single gene, it is also possible to better understand how methylation in different positions may affect the contribution of a particular gene to the aging process. Currently, however, there are only a few EPIC array data sets publicly available. Another interesting application for deep learning in the aging field is the creation of epigenetic clocks that directly predict mortality. Currently, the state-of-the-art mortality predictor is GrimAge [63], which was create based on a linear Cox proportional hazards model. We anticipate that using neural networks to include non-linear relationships and CpG-CpG interactions would result in a better lifespan predictor.

Overall, we have shown that deep learning represents an improvement in performance over current approaches for epigenetic clocks while at the same time providing new, relevant biological insights about the aging process.

## Methods

### DNA methylation data sets

For model training and testing, we gathered normal tissue samples from 143 publicly available DNA methylation data sets from the Gene Expression Omnibus, Array Express, and The Cancer Genome Atlas, comprising of the platforms Illumina Infinium Human-Methylation27 and the Illumina Infinium HumanMethylation450. All selected data sets had both processed beta values and age available for all samples. Missing values per data set were imputed with a KNN imputer from scikit-learn. Next, the data was normalized using the beta mixture quantile normalization (BMIQ) with the optimized code from Horvath, called BMIQCalibration [3, 64]. 13 data sets contained tumor samples which were separated for later analysis. Samples that failed BMIQ normalization were removed. Then, each data set was split 60% for training (n = 8999) and 40% for testing (n = 6091). The within-data set split ensures the distribution of age, gender, and tissue type between training and testing sets are unbiased (Supplementary Figure S2). In the training set, the data was further randomly subdivided by data set, with 77 (n = 4339) for model selection and 76 (n = 4660) for validation. The full list of data sets used is available in the paper’s GitHub repository (https://github.com/rsinghlab/AltumAge).

For twelve data sets in which gestational week was available, the encoding for age is the following:

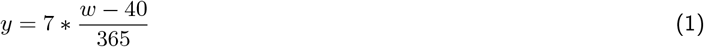

where *w* is the gestational week, and *y* is the age in years. A gestational week below 40 would have negative age; for instance, 30 weeks would be encoded as 7 * (30 – 40)/365 = −0.192. In the US in 2013, the average birth occurred at an estimated 38.5 weeks [65]. This number has changed slightly over time, and since preterm deliveries skew the average more than late-term births, we considered gestational week 40 as age 0 in such data sets.

For the cancer age acceleration analysis, we compared the test set of 13 data sets with the aforementioned separated cancer samples. These data sets were GSE32393, GSE37988, GSE26126, GSE63384, GSE59157, GSE32867, GSE30759, GSE31979, GSE77955, GSE52068, GSE49149, GSE39004. We further added GSE53051, which contains normal and tumor samples from five tissue types, for the analysis. It is worth noting that analyzing GSE53051 separately did not change the outcome of higher tumor age acceleration predicted with AltumAge vs. no difference with Horvath’s model. In total, we compared 732 normal and 3712 cancer observations across 10 different tissue types (Figure 6).

### CpG site annotation

For the annotation of CpG sites, GENCODE and Zhou et al’s annotations were used [66, 67]. 41 data sets from ENCODE with the 18-state ChromHMM information were gathered [33]: ENCFF717HFZ, ENCFF718AGZ, ENCFF371WNR, ENCFF318XQO, ENCFF340OUL, ENCFF893CAJ, ENCFF151PZS, ENCFF098CED, ENCFF273PJW, ENCFF377YFI, ENCFF773VYR, ENCFF928QES, ENCFF786HDE, ENCFF827FZN, ENCFF364PIY, ENCFF802QCI, ENCFF021NNN, ENCFF510ZEI, ENCFF175NGE, ENCFF670DBL, ENCFF825ZCZ, ENCFF912ILE, ENCFF725WBV, ENCFF829SZB, ENCFF483NRC, ENCFF717RYX, ENCFF249ZBG, ENCFF205OTD, ENCFF765OKG, ENCFF820YPQ, ENCFF685BMF, ENCFF545ZMG, ENCFF294UQS, ENCFF104ZSA, ENCFF370EGY, ENCFF860FWW, ENCFF177TTP, ENCFF151ZGD, ENCFF743GHZ, ENCFF990YHL, and ENCFF036WIO. Since AltumAge is a pan-tissue clock, the mode of each state was chosen for each CpG site.

### Model selection

Multiple machine learning models were tested in the validation set. The evaluation metrics were median absolute error (MAE), mean squared error (MSE), Pearson’s correlation coefficient (R), and median error.

To validate and train the models, the beta value of each CpG site was scaled with a robust scaler which removes the median and scales according to the interquantile range. A robust scaler was chosen as opposed to a standard scaler (mean = 0, var = 1) to better resist distortions caused by outliers. In addition, only 20,318 CpG sites common to all three platforms Illumina Infinium HumanMethylation27, Illumina Infinium HumanMethylation450, Infinium Methylation EPIC were chosen as features.

The non-neural network models trained with `scikit-learn` were support vector regression (with an RBF kernel) and random forest with the standard hyperparameters. ElasticNet, trained with `glmnet`, used the built-in λ optimization with parameters alpha = 0.5 and n_splits = 10. Moreover, for ElasticNet, Horvath’s age transformation was used [3].

To select the best performing neural network with `tensorflow`, we conducted a grid search with the following hyperparameters: number of hidden layers (2, 5, or 8), number of neurons per dense layer (32 or 64), activity and kernel regularization coefficients (0, 0.0034, or 0.0132), dropout (0 or 0.1), Gaussian dropout (0 or 0.1), batch normalization (yes or no), activation function (ReLU or SeLU), and learning rate (0.0002, 0.0005, or 0.001). The following parameters were held constant: optimizer (Adam), batch size (256), number of epochs (300), loss function (MSE), and learning rate decay by a factor of 0.2 after a 30-epoch plateau in the training loss. The weights with the lowest training loss were chosen. After selecting the best hyperparameters, we trained the neural network with adversarial regularization with `neural_structured_learning` with multiplier=0.05, adv_step_size=0.005.

We dubbed the best performing deep neural network as AltumAge. It consists of 5 hidden layers, 32 neurons per layer, activity and kernel regularization coefficients of 0.0034, no dropout, gaussian dropout of 0.1, with batch normalization, SeLU activation, and learning rate of 0.001. AltumAge was also tested in the validation set with a smaller set of features using the CpG sites selected from the ElasticNet.

To compare our deep learning approach with other state-of-the-art neural networks, we tried TabNet, an attentive interpretable tabular learning method [17]. Similarly, TabNet was trained for 300 epochs.

The results for the validation set with the full list of models, including the replication of Horvath’s model [3], is in Supplementary Table S1. Support vector regression was by far the worst performer (MAE = 14.229, MSE = 458.956), followed by random forest (MAE = 6.833, MSE = 165.354). All other models displayed R > 0.9, with AltumAge having the lowest MAE (3.563) and MSE (57.071).

### SHAP and DeepPINK

To obtain the SHAP values for AltumAge, the function `GradientExplainer` from shap was used on the test set. For the DeepPINK importance values and feature selection, the standard architecture and number of epochs was used [26]. To create the knockoff features for DeepPINK, the function `knockoff.filter` from the R 4.0.2 package `knockoff` version 0.3.3 was used with the importance statistic based on the square-root lasso.

Both SHAP and DeepPINK importance values were normalized so that their sum would equal to 100. Each importance value thus represents a percent contribution of a certain feature.

## Statistical Analysis

All statistical tests were conducted with packages `scikit-learn`, `statsmodel`, or `scipy`. Tests that were one-sided were explicitly mentioned in the main text; all others were two-sided.

To assess the performance of the models, we used median absolute error (MAE), mean squared error (MSE), Pearson’s correlation coefficient (R), and median error.

## Code Availability

Detailed instructions on how to run AltumAge can be found in the paper’s GitHub repository (https://github.com/rsinghlab/AltumAge). All code was run with Python (3.9.1) and packages `scikit-learn` (0.24.2), `pandas` (1.3.0), `numpy` (1.19.5), `glmnet` (1.1), `tensorflow` (2.5.0), `neural_structured_learning` (1.3.1), `statsmodel` (0.10.2), `scipy` (1.7.0), `shap` (0.39.0).

## Data Availability

The list of all the data sets used, a summary of the results per data set, and detailed instructions to run AltumAge can be found in the paper’s GitHub repository (https://github.com/rsinghlab/AltumAge). The GitHub also links to a Google Drive where our gathered DNA methylation data is publicly available.

## Author Contributions

L.P.L.C conceived of the presented idea. R.S and L.P.L.C designed the methodology and the experiments. L.P.L.C conducted all the experiments. L.R.L assisted with the analysis and biological interpretation of the results. All authors discussed the results and contributed to the final manuscript.

## Supplementary Information

### DeepPINK

To support the results obtained by SHAP, we also applied another method of determining feature importance called DeepPINK [26]. It works by comparing the original features with fake features. The knockoff features can be generated in many different ways, as long as they simulate the original data structure but are not related to the output. DeepPINK contrasts the relevance of the fake features against the regular input features to determine which ones are truly related to the output. It can also be used for feature selection with a controllable false discovery rate (FDR). It is worth highlighting the difficulty in feature selection in DNA methylation data. Most experiments have a couple dozen or a couple hundred samples. Depending on the type of platform used, the number of beta values for the CpG sites analyzed can vary from around 27 thousand to around 850 thousand. DeepPINK, even with a high FDR of 0.5, only selected 78 features. The fact that other sets of CpG sites unrelated to Horvath’s 353 also perform similarly well emphasizes the difficulty in finding the “true” age-related CpG sites.

#### AltumAge captures relevant age-related CpG-CpG interactions

The top 9 most important CpG sites according to SHAP account for 9.54% of total DeepPINK model importance.

#### Characterization of CpG sites by model interpretation

For the CTCF binding site analysis, the top 1000 CpG sites comprises 47.3% of DeepPINK importance. For the ChromHMM analysis, CpG importance values are also impacted by ChromHMM state for DeepPINK (Kruskal-Wallis H-test p-value = 2.982e-05). The chromatin state with the highest DeepPINK normalized median importance was heterochromatin (DeepPINK importance = 2.47e-14%, top 66th percentile of all CpG sites). Despite there being only 29 CpG sites characterized as heterochromatic, this result emphasizes the importance of chromatin packing with aging, as it is related to genome stability and maintenance.

#### Aging-related pathways

None of the CpG sites in SIRT genes appear very relevant for DeepPINK. cg21770145, located in SIRT7, accounts for 7.89e-12% of total DeepPINK importance and ranks 1342, with the highest SIRT DeepPINK importance value. For the mTOR pathway, cg05546044, located in MAPK1, has the highest DeepPINK importance of 0.029%, ranking 233. mTOR was not particularly relevant, with its most important CpG site being cg07029998 (DeepPINK importance = 1.12e-12%, rank 2459). All AMPK-related CpG sites had low (less than 10e-13%) DeepPINK importance values.

### Tables and Figures

**Table S1:**
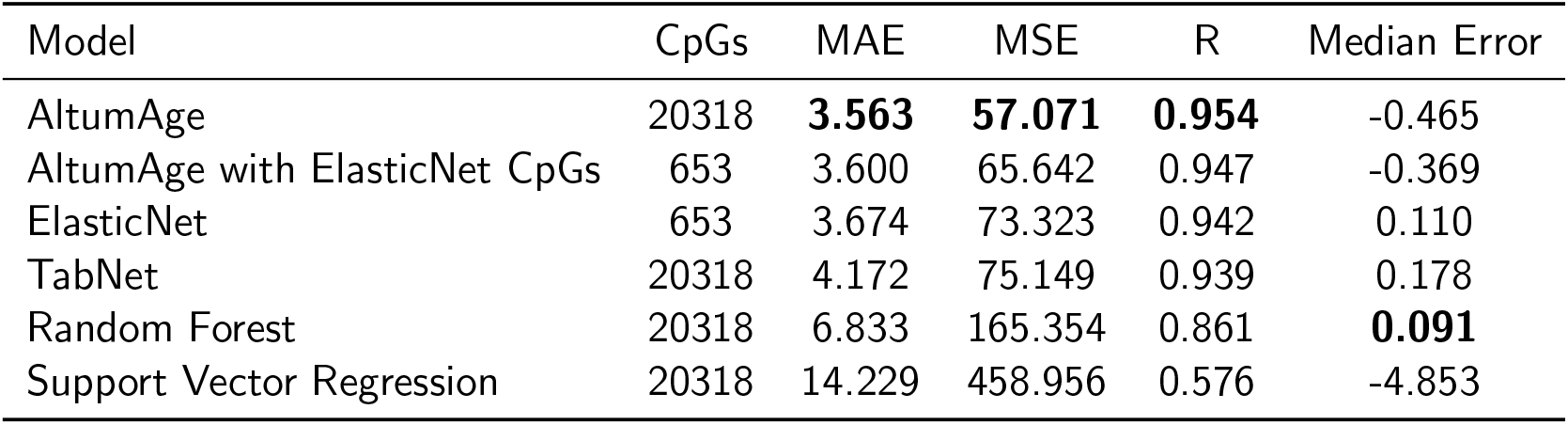
Evaluation metrics for all models in the validation set. The median absolute error (MAE) and the median error are in units of year, while the mean squared error (MSE) is in units of year-squared.

| Model | CpGs | MAE | MSE | R | Median Error |
| --- | --- | --- | --- | --- | --- |
| AltumAge | 20318 | **3.563** | **57.071** | **0.954** | -0.465 |
| AltumAge with ElasticNet CpGs | 653 | 3.600 | 65.642 | 0.947 | -0.369 |
| ElasticNet | 653 | 3.674 | 73.323 | 0.942 | 0.110 |
| TabNet | 20318 | 4.172 | 75.149 | 0.939 | 0.178 |
| Random Forest | 20318 | 6.833 | 165.354 | 0.861 | **0.091** |
| Support Vector Regression | 20318 | 14.229 | 458.956 | 0.576 | -4.853 |

**Table S2:** List of ChromHMM states by ChromHMM state ID.

| ChromHMM state ID | ChromHMM state |
| --- | --- |
| 1 | Active TSS |
| 2 | Flanking TSS |
| 3 | Flanking TSS Upstream |
| 4 | Flanking TSS Downstream |
| 5 | Strong transcription |
| 6 | Weak transcription |
| 7 | Genic enhancer1 |
| 8 | Genic enhancer2 |
| 9 | Active Enhancer 1 |
| 10 | Active Enhancer 2 |
| 11 | Weak Enhancer |
| 12 | ZNF genes and repeats |
| 13 | Heterochromatin |
| 14 | Bivalent/Poised TSS |
| 15 | Bivalent Enhancer |
| 16 | Repressed PolyComb |
| 17 | Weak Repressed PolyComb |
| 18 | Quiescent/Low |

**Figure S1:**
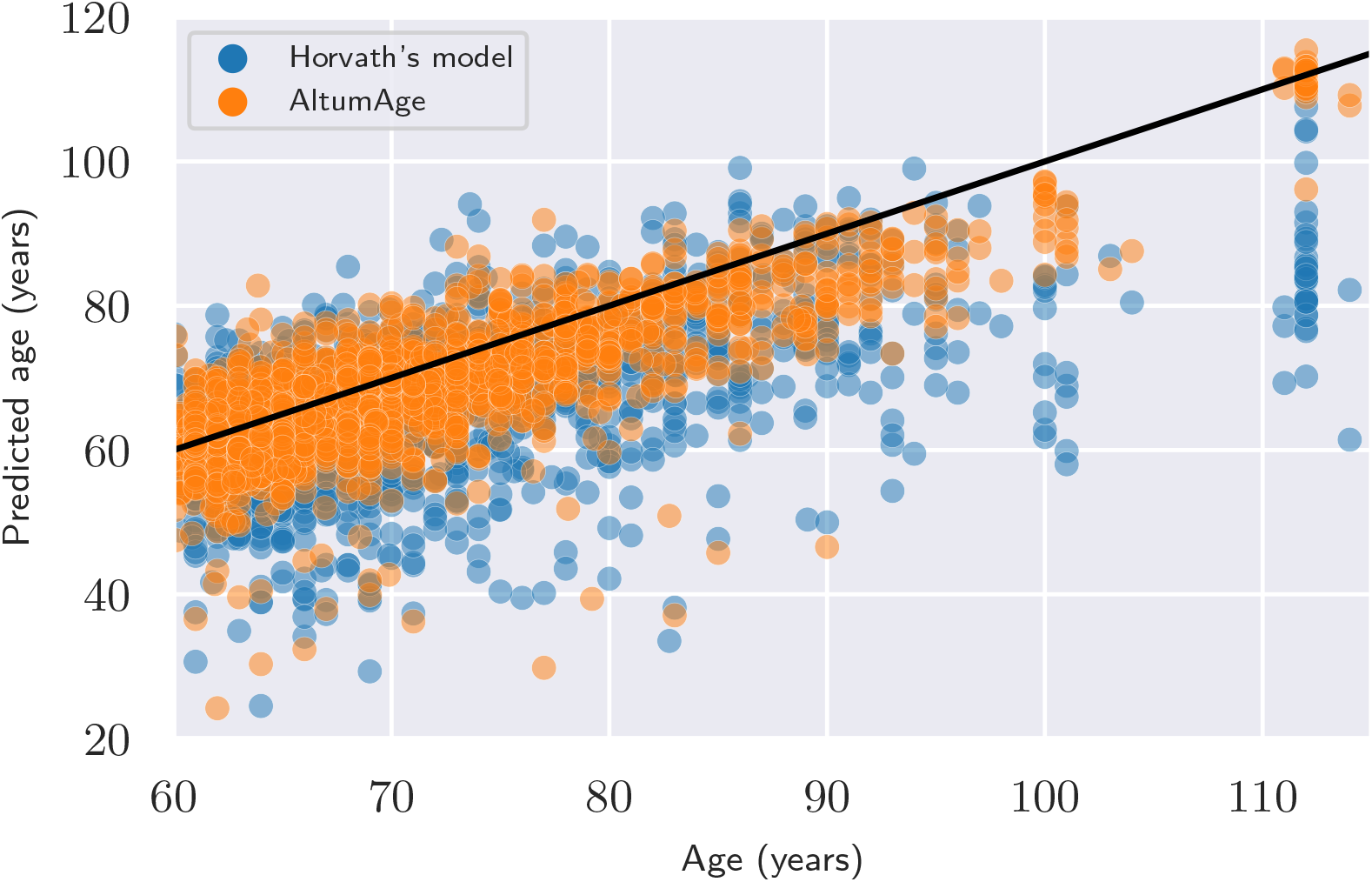
Scatter plot showing the improved performance of AltumAge in comparison to Horvath’s 2013 model for older ages. The black line represents the location where the predicted age equals the real age. AltumAge’s predictions are generally closer to the black line. Horvath’s predictions tends to give lower performance in higher ages.

**Figure S2:**
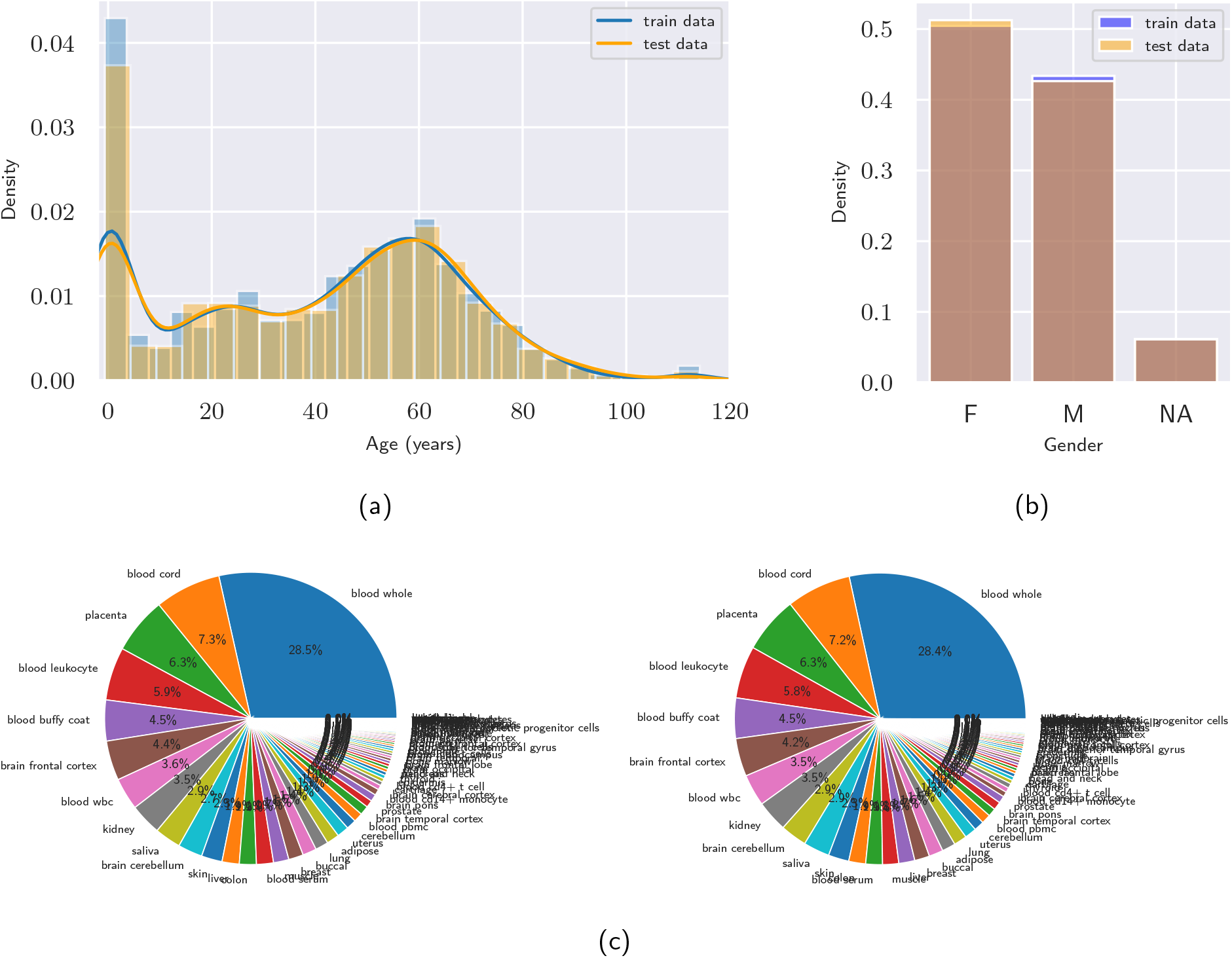
Distribution of age, gender, and tissue type in both training and testing sets. The distributions of age (a) and gender (b) are virtually identical between the two sets. Similarly with the distribution of tissue type (c) in training (left) and testing (right).

**Figure S3:**
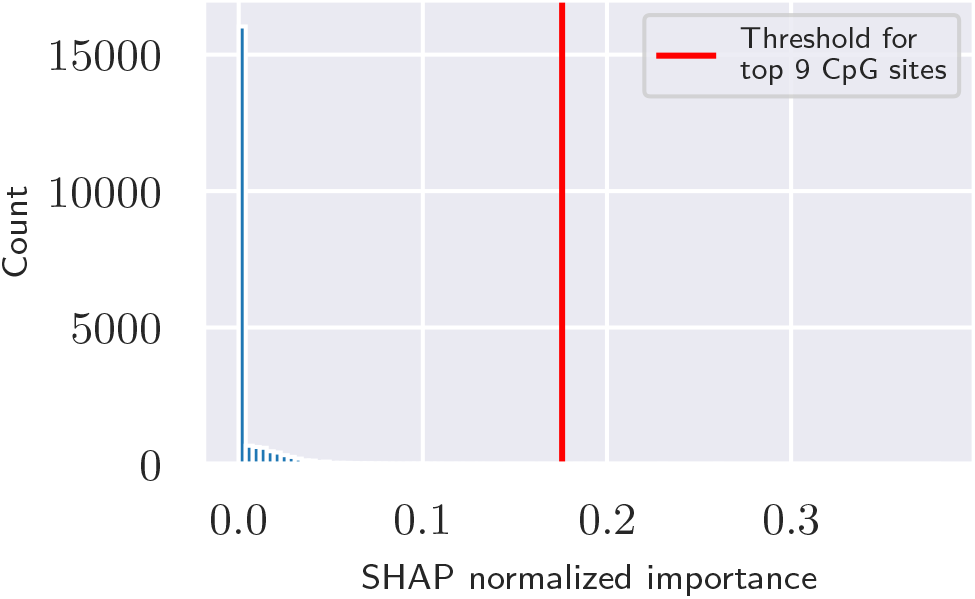
Histogram of the normalized importance values of all AltumAge CpG sites according to SHAP. The red line represents the threshold for the top nine CpG sites. These have a much higher importance than most other CpG sites.

**Figure S4:**
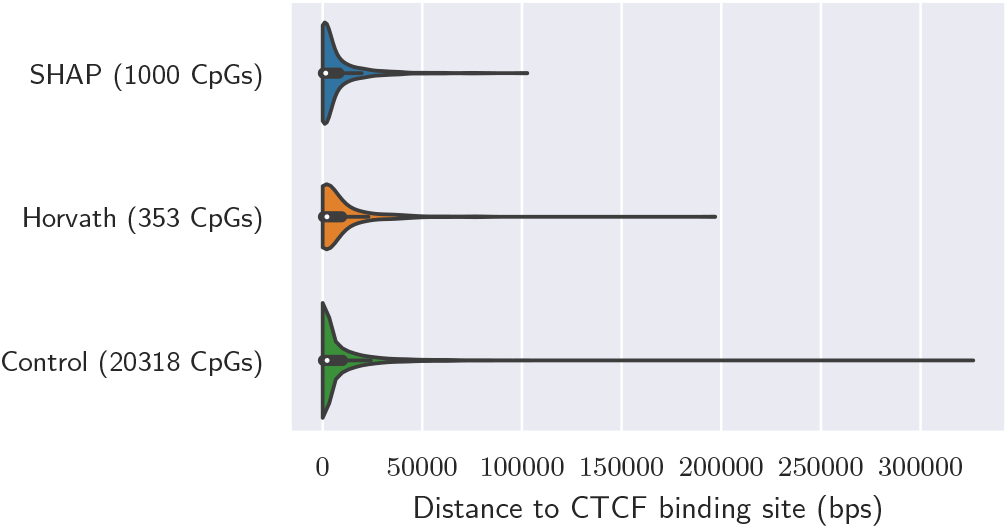
Violin plot showing that the top 1000 CpG sites according to SHAP are closer to CTCF binding sites than the 20,318 control CpG sites. Horvath’s CpG sites are not statistically significantly different from the control.

**Figure S5:**
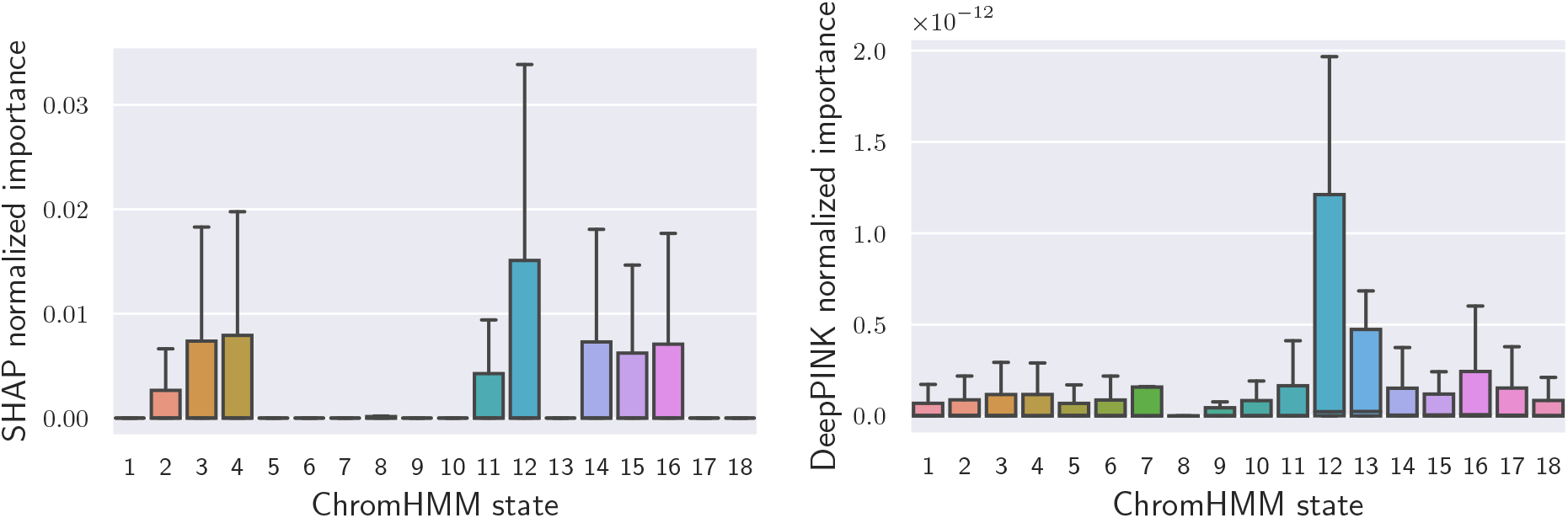
Box plots of SHAP and DeepPINK normalized importance values by ChromHMM state. Outliers were removed for better figure visualization. No specific ChromHMM state stands out in importance.

**Figure S6:**
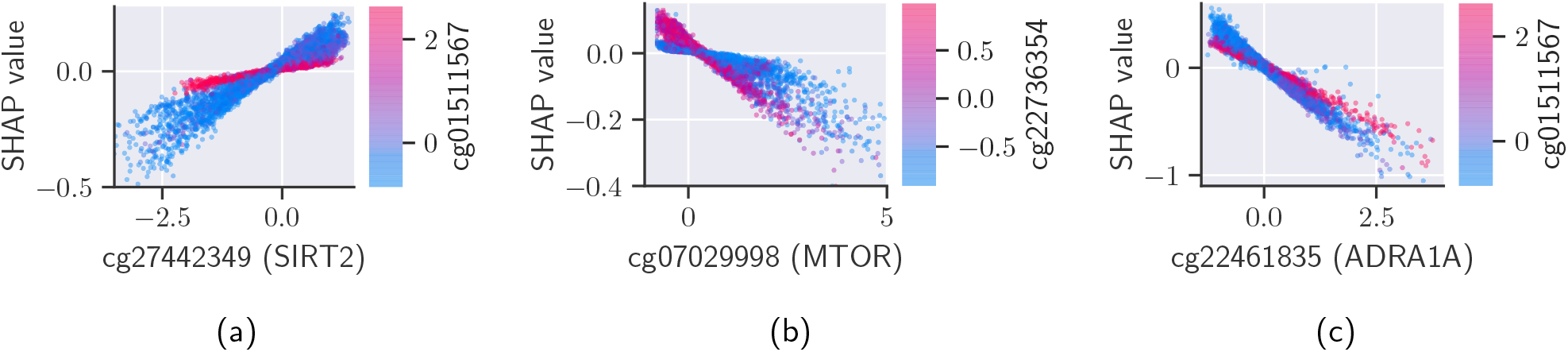
SHAP dependence plots of three CpG sites in SIRT2, mTOR, and ADRA1A. The x-axis shows the standardized beta values for each specific CpG site; the y-axis, its SHAP value, and the coloring scheme, the scaled beta values for the CpG site with the highest interaction. These are the most important CpG sites according to SHAP for AltumAge in the SIRT and mTOR pathways.

**Figure S7:**
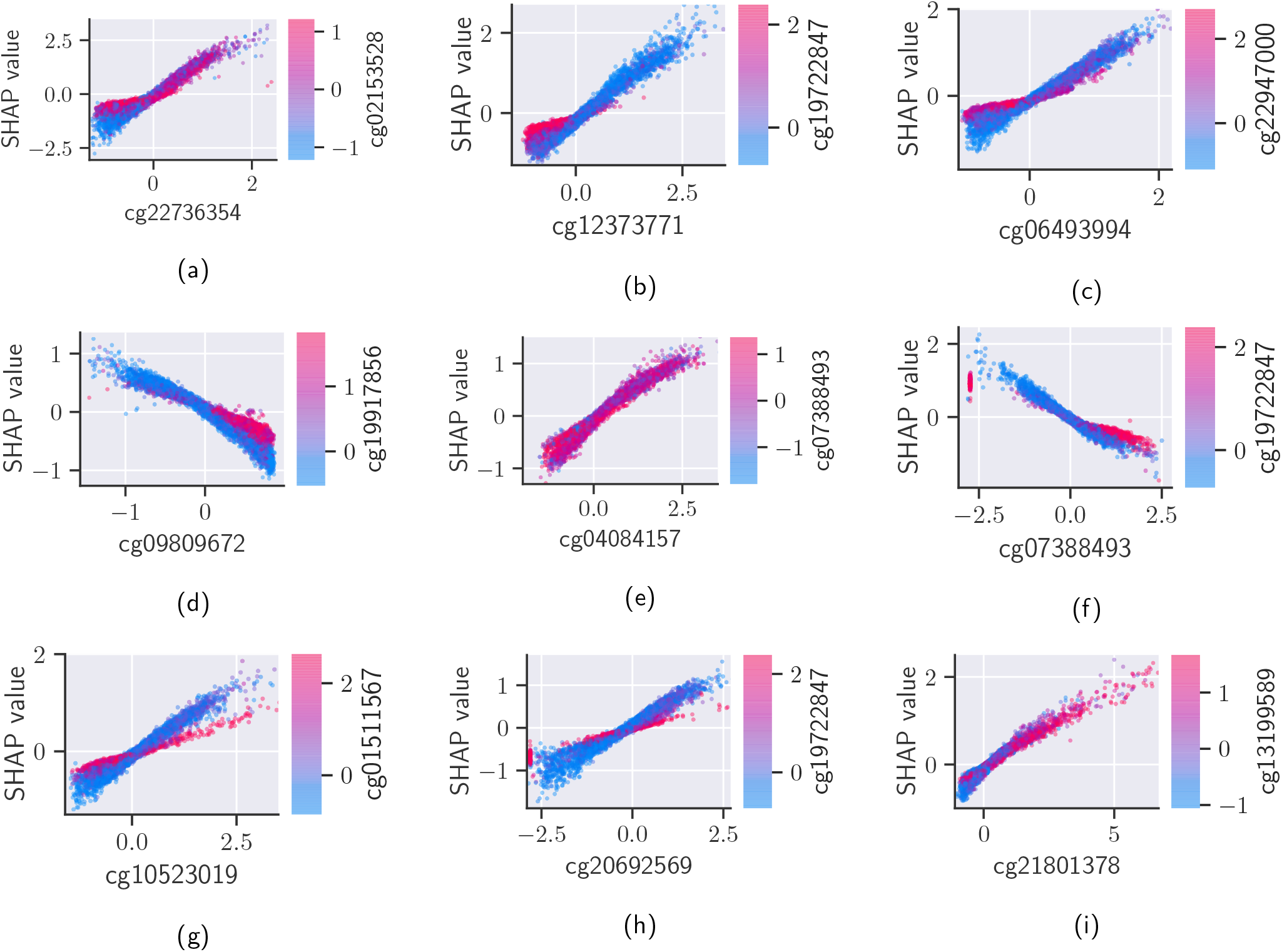
Dependence plots of the nine most important CpG sites (a-i) in AltumAge based on SHAP values. They are ordered from top left to bottom right in terms of importance. The x-axis shows the scaled beta values for each specific CpG site; the y-axis, its SHAP value, and the coloring scheme, the scaled beta values for the CpG site with the highest interaction. The effect of a specific CpG site on the predicted age can vary based on a secondary CpG site.

**Figure S8:**
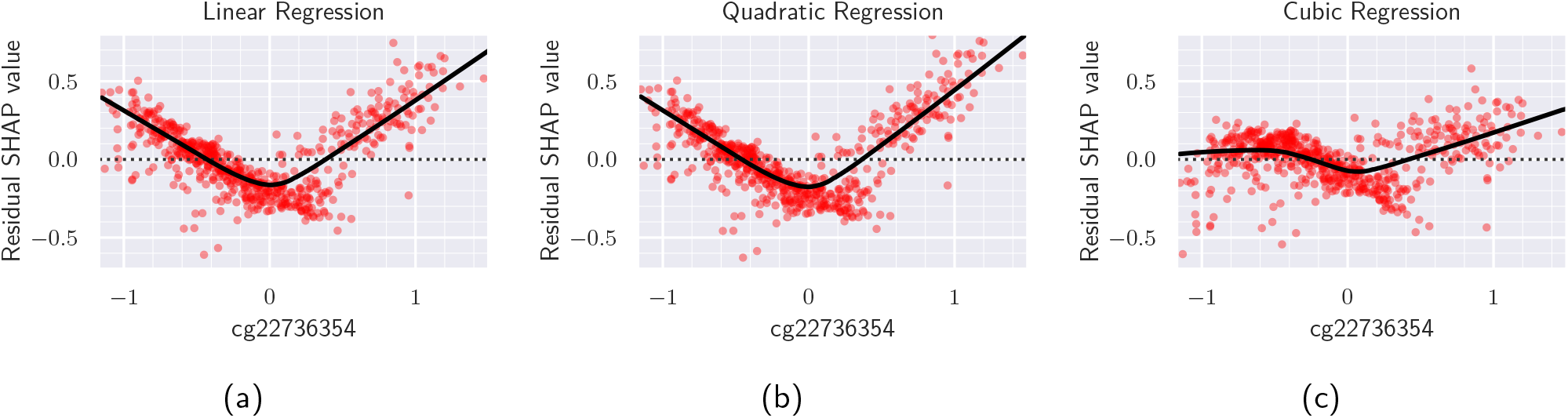
Residual plots of cg22736354 by order of regression. A linear regression (a) underestimates around the boundaries and underestimates in the middle, demonstrating the non-linear relationship. The same occurs with a quadratic regression (b). When the order is increased to three, the cubic regression (c) takes the non-linearity better into account.

**Figure S9:**
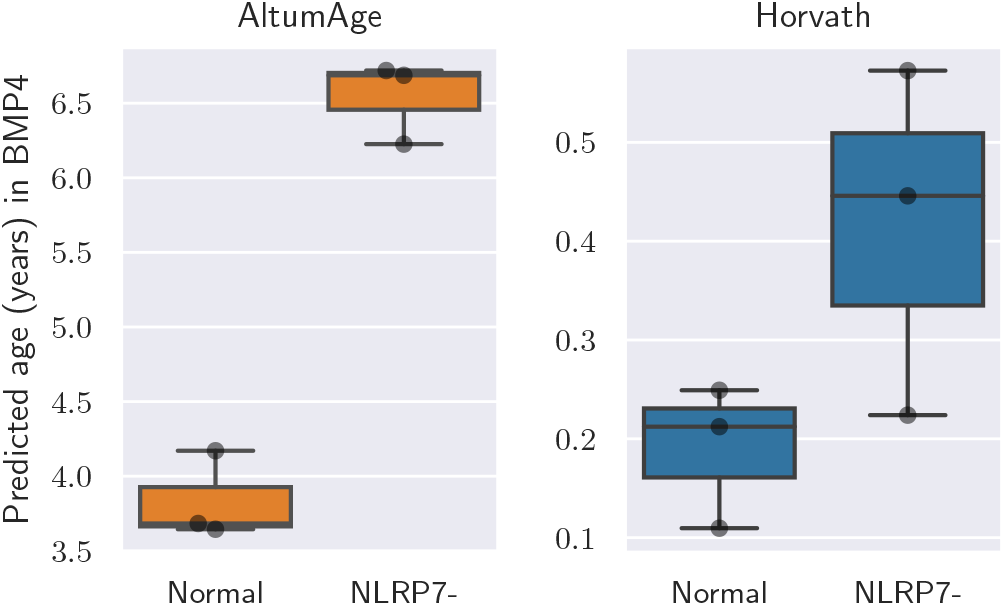
Box plots showing predicted age of H9 ESCs with NLRP7 knockdown (NLRP7-) or control (Normal) in BMP4 differentiating medium.

## Footnotes

1 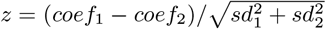, here *coef* is the coefficient, *sd* is the standard deviation

